# Anticipatory modulation of motor unit discharge rate before rapid isometric elbow flexion force production

**DOI:** 10.64898/2026.07.11.737916

**Authors:** Ji-Won Park, Sungjune Lee, Yoon-Seok Choi, Dawon Park, Hyeon Hur, Hyun-Soo Kim, Jaeho Park, Jaebum Park

## Abstract

Surface electromyography (EMG) studies have demonstrated anticipatory muscle activation prior to predictable voluntary actions. However, the mechanism through which this preparation is expressed, whether through motor-unit recruitment or discharge-rate modulation, remains to be elucidated. We employed surface decomposition EMG to quantify motor-unit behavior prior to self-paced rapid isometric elbow-flexion force pulses. Twelve healthy young men were instructed to generate elbow-flexion pulse force at 30%, 40%, and 50% of the maximal voluntary contraction (MVC) at a self-selected time. Motor-unit activity was decomposed from the biceps brachii and triceps brachii muscles, and the normalized active motor-unit number and mean discharge rate were analyzed prior to pulse onset. In the agonist, the pre-pulse increase mean discharge rate exhibited a higher value than the change in detected motor-unit count, particularly at the 40% and 50% MVC targets. The discharge-rate increase scaled with the target force, with a heightened response observed in high-threshold as compared to low-threshold motor units. Antagonist recordings with sufficient decomposition yield exhibited a similar discharge-rate-dominant pattern; however, these data were available from a smaller sample size. Present findings suggest that discharge-rate modulation is a primary motor-unit-level feature of anticipatory preparation for rapid isometric force production.

## 1. Introduction

Anticipatory adjustments are defined as feedforward changes in muscle activation and force that occur prior to predictable voluntary actions. These adjustments are intended to prepare the neuromuscular system for the mechanical consequences of an upcoming action while limiting premature changes in task-level output. In tasks involving opposing muscles, preparatory agonist–antagonist activation may occur with minimal change in net force because their mechanical effects partially counterbalance one another. This adjustment is a critical function for the sequence of the purposeful destabilizing followed by recovering the stability of a particular movement in humans (Zhou et al., 2013). A substantial body of research has been dedicated to substantiating the hypothesis that humans employ an anticipatory adjustment, that is, a feedforward mechanism, whereby subtle alterations in activation levels occur before initiation of the intended action. This encompasses all involved elements, including changes in net mechanical outcomes produced by multiple effectors. Anticipatory adjustments may therefore be expressed at the level of force, muscle activation, or multi-effector coordination. In certain tasks, preparatory activation of agonist and antagonist muscles can occur with minimal change in net mechanical output because the opposing muscle actions partially counterbalance one another (Kong et al., 2019; Song et al., 2023). A prerequisite for feedforward changes is that the performer has advance knowledge of upcoming actions, such as time, direction, and magnitude, which are associated with the properties of the upcoming mechanics against predictable perturbations (Krishnan et al., 2012).

The majority of research on anticipatory adjustment has centered on changes in covariation between multi-finger force production (Olafsdottir et al., 2005; Shim et al., 2005; Park and Xu, 2017) and multi-muscle activation during postural adjustment (Klous et al., 2011; Krishnan et al., 2011; Kaewmanee et al., 2022). These studies have provided critical information regarding the strategies of the module configuration (Torres-Oviedo & Ting, 2010; Roh et al., 2012) or covariation patterns of multiple units of the grouped muscles (i.e., M-mode, Krishnamoorthy et al., 2003; Asaka et al., 2008). In the context of anticipatory strategies within the muscular system, the investigation of activation patterns across multiple motor units emerges as a pivotal area of inquiry. The recruitment of these motor units determines their availability for anticipatory responses, while discharge-rate coding determines the strength and rapidity of their action. The integration of spatial and temporal patterns within the nervous system facilitates the stabilization of the body, the preparation of relevant joints, the prevention of premature movement, and the precise generation of intended actions. However, the conventional surface electromyography (EMG) system was unable to discern whether anticipatory muscle activation signifies the recruitment of additional motor units, an escalation in the discharge rate of already active units, or a combination of both. Recently, the surface decomposition EMG systems facilitate investigation of this question. In particular, muscle force is derived from the integration of two distinct functions of motor-unit activation, namely the number and firing frequency of the detected active motor units. The integration of these factors yields the following expression for a particular muscle force: muscle force ≈ ∑ *f_1_*(F_MU_)·*f_2_*( _MU_), where **F** represents force by individual motor units, D denotes the discharge frequency of the corresponding motor unit, and *f_1_* and *f_2_* represent the functions of the detected active motor unit number and mean discharge rate, i.e., frequency, respectively (Bawa and Murnaghan, 2009). The size principle (Henneman, 1957) has been shown to be a reliable predictor of which units are most likely to become operational at the earliest stages of a contradiction. However, the precise magnitude and velocity of the anticipatory response are determined by a complex interplay of various factors including the recruitment number, the unit force capacity, the recruitment timing, and the instantaneous discharge rate. Consequently, a multitude of combinations of number and firing frequency could yield analogous force outcomes, thereby aligning with the concept of motor redundancy (Bernstein, 1967; Latash et al., 2002).

As demonstrated in prior experimental findings, firing frequency has been identified as a major contributor for the initiation of muscle activation, leading to the production of pulse force (De Luca & Contessa, 2012). This phenomenon occurs subsequent to low-level force production. Therefore, it is reasonable to hypothesize that the primary factor contributing to changes in muscle activation during the anticipatory process is motor-unit firing frequency. In this study, anticipatory change at the motor-unit level was operationally defined as advanced motor-unit modulation prior to detectable changes in task-level variables, such as net force or torque.

To the best of our knowledge, no study has examined whether anticipatory motor-unit regulation is expressed primarily through alterations in the number of detected active motor units or through modifications in their discharge rate. In the present experiment, we investigated whether anticipatory motor-unit modulation before self-paced rapid elbow-flexion force production is dominated by discharge-rate modulation rather than recruitment-number modulation. The target muscles selected for this study were the biceps brachii and its antagonist, the triceps brachii, which are involved in elbow flexion and extension.

## 2. Methods

### 2.1. Participants

Twelve right-handed young men (25.21 ± 2.78 years, 174.43 ± 5.54 cm, 73.64 ± 6.89 kg) participated in this experiment. Hand dominance was determined using the Edinburgh handedness inventory (Oldfield, 1971). Given the exclusive inclusion of young males in the present study, it is imperative to exercise caution when generalizing the findings to other demographic groups, such as women, older adults, or clinical populations, until further research is conducted. The study was approved by the Institutional Review Board of Seoul National University (IRB No. 2101/002-001) and was conducted in accordance with the Declaration of Helsinki. Prior to participation, all participants provided written informed consent, and the principles of their privacy rights were observed.

### 2.2. Apparatus

A single strain gauge transducer (MC3A-1000, AMTI, USA) was used to measure force, with a sampling frequency of 200 Hz. A wireless decomposition electromyography (dEMG) system (Trigno Galileo sensor, Delsys, USA) was used to record an array of differential surface EMG signals. These signals were sampled at 2000 Hz with a 20–450 Hz band-pass filter. The dEMG electrodes were positioned on the belly of the biceps brachii and the lateral head of the triceps brachii in accordance with SENIAM guidelines (Hermens et al., 2000). Participants were seated on a chair, with a 27-inch (68.58 cm) monitor positioned 1 m away at eye level. The shoulder, trunk, and upper arm were configured as shoulder flexion and upper arm abduction about 45°, and this configuration was fixed; thus, the motion of these segments was restricted during the task. The elbow joint was flexed 90° in the sagittal plane, and the forearm was supinated and placed on a customized concave-shaped pad to prevent extension movement (Fig. 1A).

**Fig. 1.**
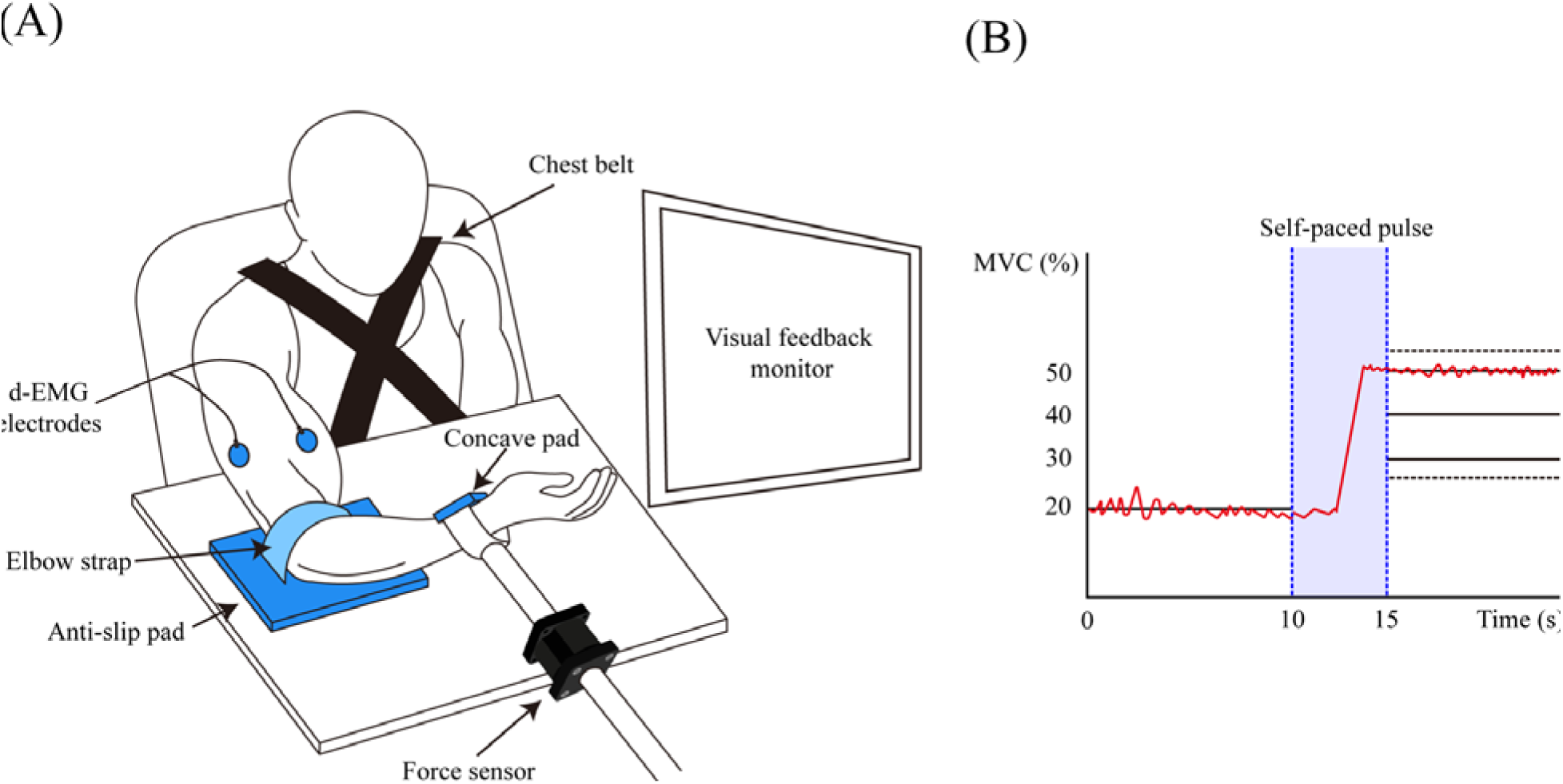
Experimental setup. Participants performed the isometric elbow-flexion task requiring quick pulse force generation **(A)**. The tested arm was supported on an anti-slip pad, and the supinated wrist was secured on a customized concave-shaped pad. During force generation, a chest belt and elbow strap were used to prevent trunk movement and stabilize the arm position, respectively. dEMG electrodes were positioned over the biceps brachii and lateral head of the triceps brachii. Visual feedback displayed the initial steady-state force level (MVC_20_) and the target force levels (MVC_30,_ MVC_40_, MVC_50_), with a 5% error margin **(B)**. Participants generated a self-paced pulse force toward the target force within the 5 s interval from 10 to 15 s.

### 2.3. Experimental procedures

The experiment consisted of one auxiliary task, maximal voluntary contraction (MVC) force production, and a main task for generating quick pulse force during isometric elbow flexion. For the MVC task, participants were instructed to produce maximum isometric elbow flexion force within 5 s. Following three attempts, the highest value was used to normalize target force level in the main task. The main task consisted of steady-state force production followed by quick pulse force during isometric elbow flexion. In a single trial, force feedback was displayed on a computer screen with two horizontal lines representing the initial steady-state force level (20% MVC) and target levels of 30% (MVC_30_), 40% (MVC_40_), and 50% (MVC_50_) of MVC, with a 5% margin of force error (Fig. 1B). Participants were required to sustain 20% MVC for 10 s and to exert quick pulse force to the target at a self-selected time within the subsequent 5 s (Fig. 1B). Participants performed 10 trials for each condition, with rest intervals to minimize fatigue. Rest intervals were 2 min between trials and 5 min between conditions.

### 2.4. Data analysis

#### 2.4.1. Curation of phases and temporal parameters

The pulse force onset time (t_0_) was defined as the time at which the first derivative of force (dF/dt) reached 5% of its maximum value (Park et al., 2012). The t_0_ variable was used as the reference time for subsequent time variables. The onset time of the anticipatory phase (t_A_) was defined as the time at which the regulation ratio of the detected active motor-unit number (R_N_) and firing frequency (R_F_) began to increase by more than two standard deviations of the average regulation ratio over the steady-state period. The peak pulse time (t_PN_ & t_PF_) was defined as the time of the maximum value of each of R_N_ and R_F_ after t_0_. Accordingly, temporal parameters (t*_j_*) of the time-series dEMG data were categorized into three phases: the steady-state phase (SP), the anticipatory phase (AP), and the pulse phase (PP). The SP (t_SP_), AP (t_AP_), and PP (t_PNP_ & t_PFP_) were established within the ranges of −1000 to −500 ms from t_0_, t_A_ to t_0_, and t_0_ to t_P_, respectively.

#### 2.4.2. Motor-unit decomposition and regulation: R_F_ and R_N_

Component motor unit action potential (MUAP) waveforms and firing times were extracted from multichannel surface EMG signals using decomposition algorithms (NeuroMap software, Delsys Inc.; De Luca et al., 2015). Motor units were retained if they exceeded the manufacturer-recommended criterion for decomposition quality and exhibited physiologically plausible discharge patterns. The number of accepted units is reported in Table 1. When a specific unit was identified across multiple trials, unit identity was confirmed using waveform similarity and discharge behavior. Baseline values of recruited MU number (M_NUM-SP_) and firing frequency (M_FREQ-SP_) were calculated as mean values during the steady-state period, from −1000 to −500 ms relative to t_0_, where the produced force was well matched to the target force level. Motor-unit regulation indices for recruited number (R_N_) and firing frequency (R_F_) were calculated using Eq. (1). The detailed procedure was as follows: (1) the spike-time train and firing frequency over time of individual motor units were obtained; (2) the number of fired motor units within the first 200 ms window was counted (M_NUM-base_); (3) firing frequency of the counted motor units was averaged (M_FREQ-base_); (4) the temporal position of the window was moved by 1 ms from −2000 to 800 ms with respect to t_0_; and (5) the time series of M_NUM_ and M_FREQ_ were first normalized to their respective baseline values M_NUM-SP_ and M_FREQ-SP_ (Eq. 1), and then further normalized to the initial value of each trial. The resulting time series were denoted normalized R_N_ and R_F_, respectively.

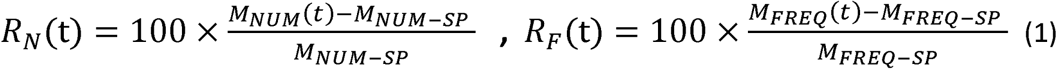

**Table 1.**
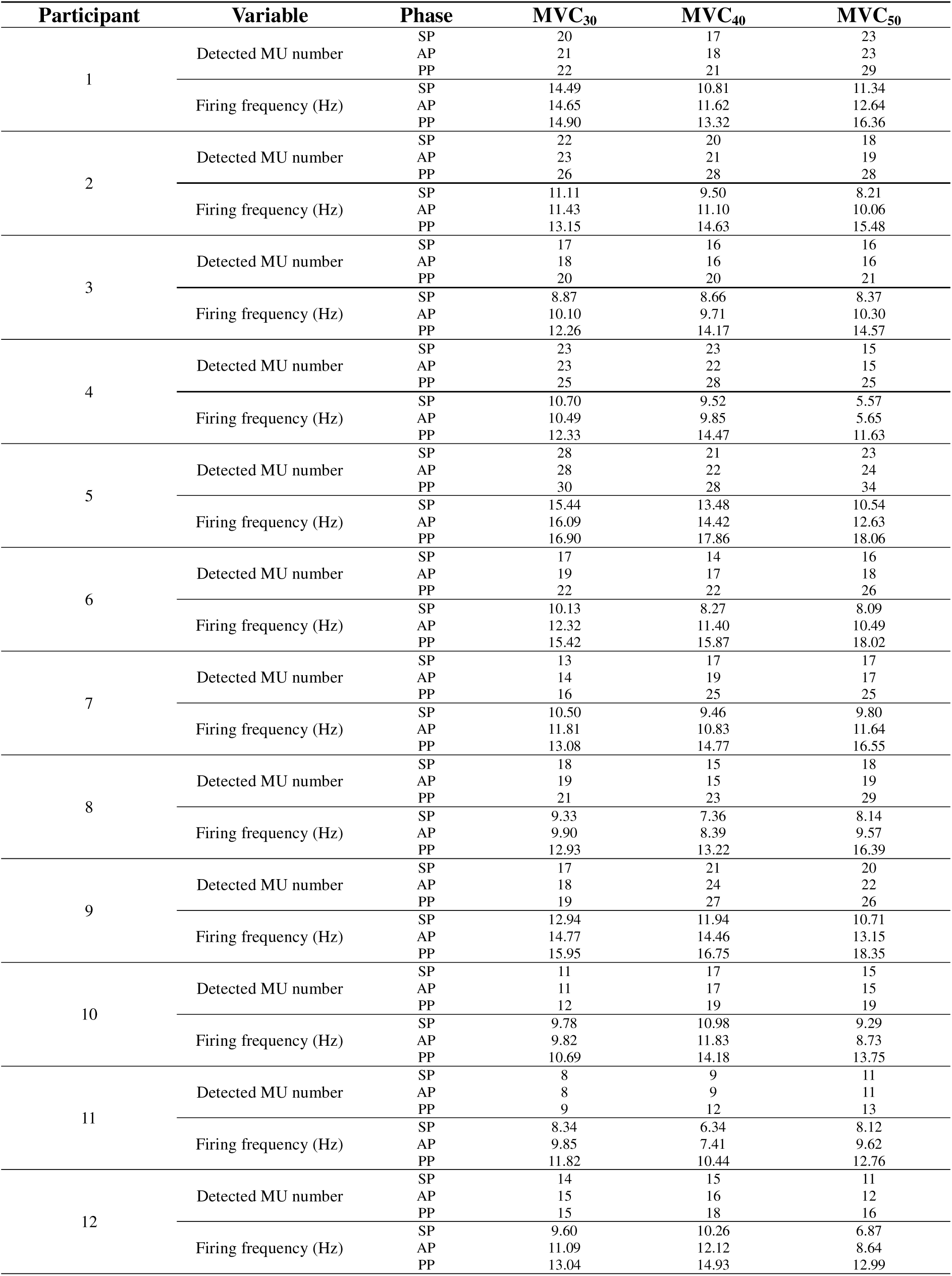
Detected motor unit number and firing frequency of the agonist muscle across phases in 12 participants.

#### 2.4.3. Indices of changes in motor-unit regulation: **Δ**R_F_ and **Δ**R_N_

The alterations in motor-unit regulation of number, ΔR_N_(t), and firing frequency, ΔR_F_(t), over time were calculated separately with respect to the time parameters t_SP_, t_AP_, and t_PP_. To conduct a sensitivity analysis, the primary comparison between ΔR_F_ and ΔR_N_ was repeated using a fixed −300 to 0 ms window before t_0_. Alterations in motor-unit regulation were quantified as the mean gradient of each phase, as described by Eq. (2). Because R_N_ and R_F_ were expressed as percentages of steady-state values, the units of ΔR_F_ and ΔR_N_ were %/s.

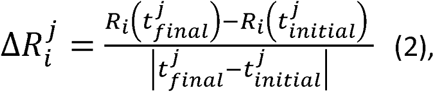

where *i* = {N, F} and *j* = {SP, AP, PP}. For AP, the initial and final times were t_A_ and t_0_, respectively. Because R_N_ and R_F_ were expressed as percentages of steady-state values, the rates of change of R_N_ and R_F_, ΔR_N_ and ΔR_F_, were expressed as %/s.

#### 2.4.4. Individual motor-unit activation: classification by recruitment threshold

The initial step entailed identification of all the detected active motor units over the entire time series of a given trial. A total of six motor units were selected for analysis from the pool of decomposed motor units. These motor units were identified based on recruitment-threshold information for individual motor units. Motor units were then ranked according to recruitment order within each trial. In accordance with the size principle, the first three recruited units were designated lower-recruitment-threshold motor units (LMU_1_, LMU_2_, and LMU_3_). The final three were classified as higher-recruitment-threshold motor units (HMU_1_, HMU_2_, and HMU_3_). The LMU/HMU firing-frequency analysis was conducted exclusively on active units that were active before pulse onset and provided sufficient discharges during AP. Firing frequencies of LMU and HMU were quantified using the inter-spike interval, the instantaneous discharge rate was calculated separately for each unit from its inter-spike intervals and then averaged within the lower-(LMU) and higher-recruitment-threshold (HMU) groups.

### 2.5. Statistical analysis

All data are presented as mean ± SD unless otherwise stated. For each participant and condition, dependent variables were averaged across accepted trials before statistical analysis; therefore, the participant was the statistical unit. Differences between pre- and post-MVC values were analyzed by a paired t-test. The three-way repeated-measures ANOVAs with factors *Force* (3 levels: MVC_30_, MVC_40_, and MVC_50_), *Phase* (3 levels: SP, AP and PP), and *Function* (2 levels: Num and Freq). Additionally, two-way repeated-measures ANOVAs were performed: one examining *Phase* (3 levels: SP, AP and PP) and *Function* (2 levels: Num and Freq), and the other examining *Force* (3 levels: MVC_30_, MVC_40_, and MVC_50_) and *Threshold* (2 levels: Low and High). Factors were selected according to the specific statistical test. Mauchly’s sphericity test was used to confirm assumptions of sphericity, and the Greenhouse-Geisser correction was used when the sphericity assumption was violated. For post hoc testing, multiple pairwise comparisons with Bonferroni correction were conducted.

Finally, the statistical power for all comparisons was estimated, thereby confirming that the power of all planned comparisons was approximately 0.7 for the pool of 12 participants. Furthermore, partial eta squared (*ηp^2^*) was reported as the effect size for all analysis of variance (ANOVA) tests. The interpretation of effect sizes was conducted in accordance with established criteria. Specifically, according to Richardson (2011), partial eta squared values ranging from 0.01 to 0.14 were classified as small, medium, and large, respectively. The level of significance for all statistical tests was set at *p* < 0.05. Analyses were carried out using IBM SPSS Statistics 26.0 (IBM, Armonk, NY, USA).

## 3. Results

Pre- and post-experiment MVC forces of the elbow joint were compared, and the results indicated no significant difference between conditions in MVC values (*p* = 0.11). Furthermore, the R_F_-dominant pattern was sustained in the fixed-window analysis from −300 ms to t_0_, with ΔR_F_ exceeding ΔR_N_ across all force levels and increasing with target force.

### 3.1. Regulation of firing rate and recruited number in the agonist muscle

During SP, R_F,_ and R_N_ in the agonist were nearly unchanged and not statistically different between MVC conditions (Figs. 2A–D, Table 1). However, the changes in the two variables, ΔR_F_ and ΔR_N_, were non-parallel in AP (Fig. 2E). Specifically, during the AP, agonist ΔR_F_ escalated progressively alongside target force, whereas ΔR_N_ did not show the same scaling pattern, leading ΔR_F_ to exceed ΔR_N_, particularly at MVC_40_ and MVC_50_. In contrast, this pattern was reversed during the PP; although both ΔR_F_ and ΔR_N_ increased progressively with target force, ΔR_N_ surpassed ΔR_F_, especially under MVC_40_ and MVC_50_. These results were supported by a three-way repeated-measures ANOVA on ΔR*_i_* with factors *Function* (2 levels: Num and Freq), *Phase* (3 levels: SP, AP and PP), and *Force* (3 levels: MVC_30_, MVC_40_, and MVC_50_), which showed main effects of *Phase* (*F*_(1.11, 12.25)_ = 62.19, *p* < 0.001, *ηp*^2^ = 0.85) and *Force* (*F*_(2, 22)_ = 25.24, *p* < 0.001, *ηp*^2^ = 0.70) with significant two-way (*Function* × *Phase*: *F*_(1.17, 12.86)_ = 38.04, *p* < 0.001, *ηp*^2^ = 0.78; *Phase* × *Force*: *F*_(1.67, 18.38)_ = 11.52, *p* = 0.001, *ηp*^2^ = 0.51) and three-way interactions (*F*_(2, 22)_ = 18.01, *p* < 0.001, *ηp*^2^ = 0.62).

**Fig. 2.**
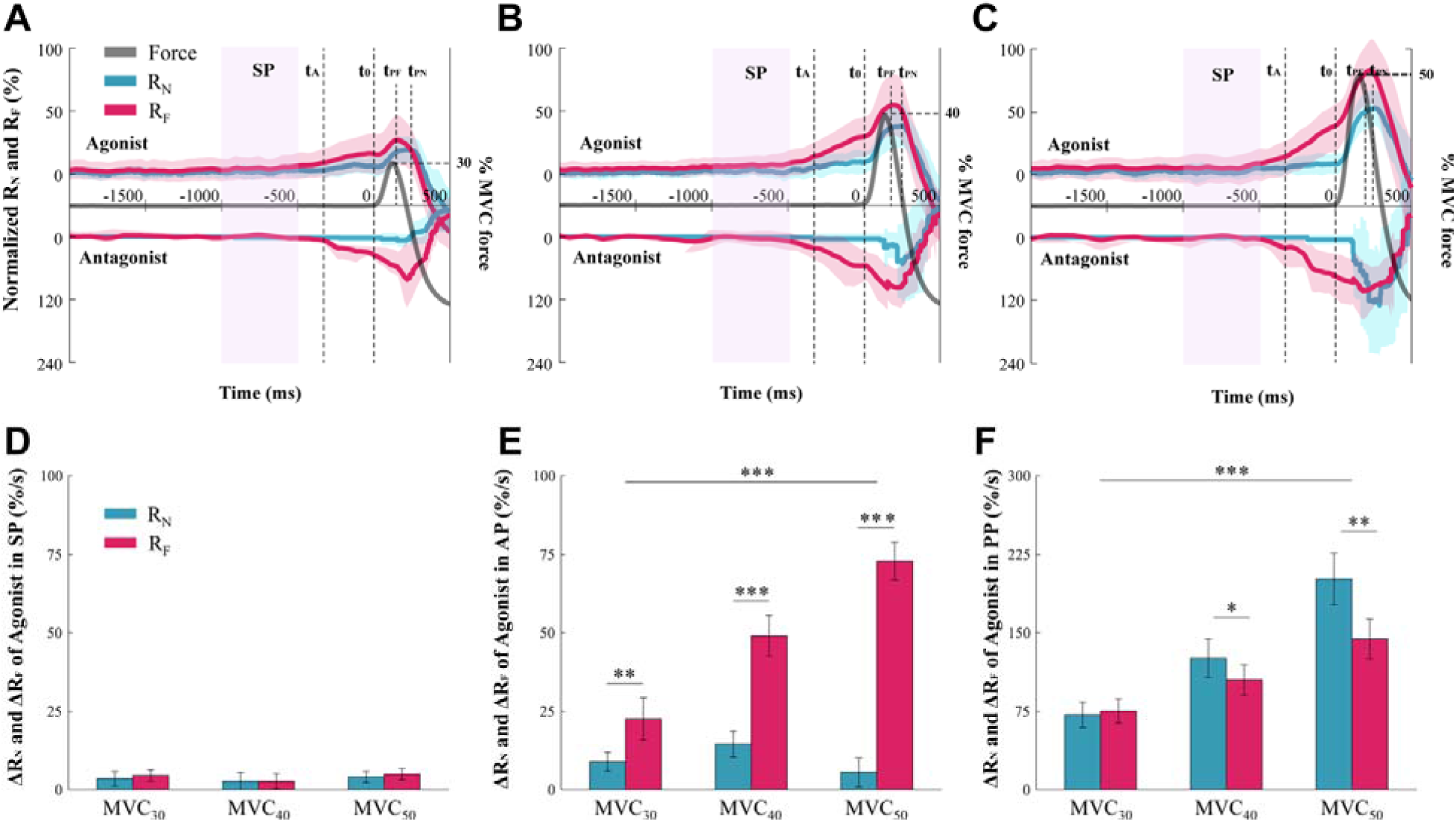
Averaged force and motor-unit regulation profiles across participants for the 30% MVC (MVC_30_; **A**), 40% MVC (MVC_40_; **B**), and 50% MVC (MVC_50_; **C**) conditions. The gray, cyan, and magenta traces indicate pulse force, normalized the detected active motor-unit number (R_N_), and normalized firing frequency (R_F_), respectively. The upper and lower profiles represent agonist and antagonist, respectively. The shaded area denotes the steady-state phase (SP). Black vertical dashed lines indicate anticipatory-phase onset (t_A_) and pulse force onset (t_0_). The magenta and cyan dashed arrows indicate the time of peak R_F_ (t_PF_) and time of peak R_N_ (t_PN_) for the agonist. Panels **D–F** compare ΔR_N_ and ΔR_F_ in agonist motor-unit regulation across force levels (MVC_30_, MVC_40_, and MVC_50_) during the steady-state phase (SP; **D**), anticipatory phase (AP; **E**), and pulse phase (PP, **F**). For each phase, ΔR_N_ and ΔR_F_ represent the rate of change, computed as the difference between the final and initial values divided by the duration of that phase. Asterisks indicate statistically significant difference: * *p* < 0.05, ** *p* < 0.01, *** *p* < 0.001

### 3.2. Firing frequency and recruitment-number changes in the antagonist muscle

The number of decomposed motor units in the antagonist was limited during SP, and large inter-subject variability was observed in both ΔR_N_ and ΔR_F_ during PP. Therefore, antagonist analysis was limited to AP. As in the agonist, changes in ΔR_N_ and ΔR_F_ were unequal (ΔR_N_ < ΔR_F_, *p* < 0.001), and these changes increased with force level, as confirmed by a *Function* × *Force* interaction (*F*_(1.30, 7.81)_ = 7.73, *p* = 0.02, *ηp*^2^ = 0.56; Fig. 4B).

### 3.3. Anticipatory changes in lower- and higher-recruitment-threshold motor units

For the agonist muscle (i.e., the biceps brachii), the changes in firing frequency of both LMU and HMU increased during AP (*Force*: *F*_(2, 22)_ = 5.05, *p* < 0.05, *ηp*^2^ = 0.32), while the relatively large increase in HMU was confirmed by a significant *Threshold* × *Force* interaction (*F*_(2, 22)_ = 4.04, *p* < 0.05, *ηp*^2^ = 0.27). In addition, post hoc comparisons confirmed that the significant effect of *Threshold* (LMU < HMU) was larger with the force levels, MVC_30_ < MVC_40_ < MVC_50_ (*p* < 0.001) (Figs. 3 & 4A, Table 2). For the antagonist muscle, i.e., the triceps brachii, the patterns of change, i.e., R_F_-dominant, were analogous to those of the agonist muscle, particularly during the AP (Figs. 2A–C). Nevertheless, the sample size for the successful antagonist decomposition was limited (*n* = 7) due to statistical power constraints. Consequently, the results of the antagonist muscle were considered tentative.

**Fig. 3.**
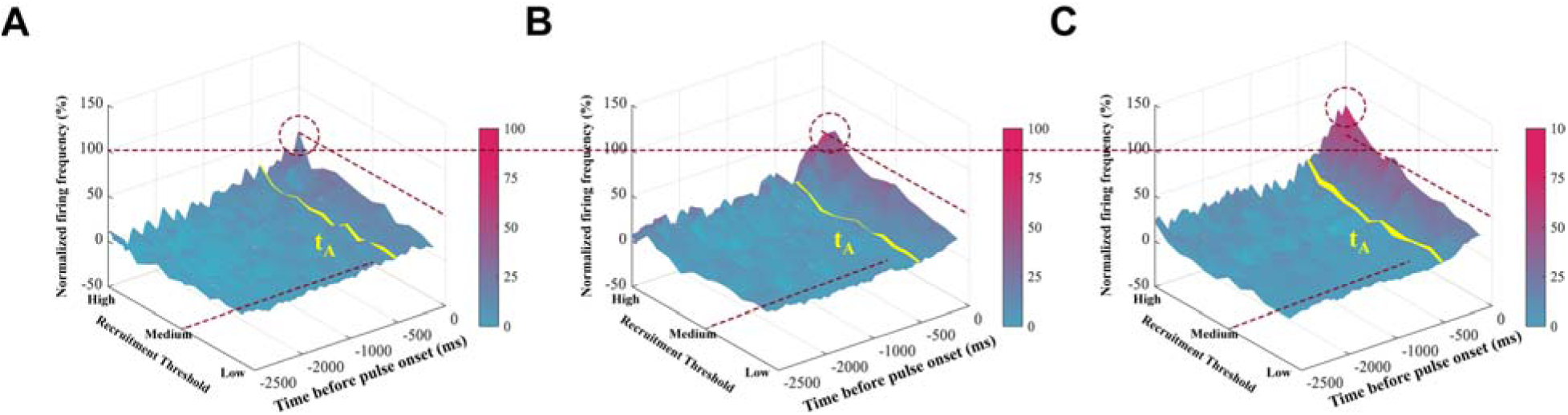
Individual agonist motor-unit firing-frequency profiles across recruitment thresholds in a representative participant for the MVC_30_ **(A)**, MVC_40_ **(B)**, and MVC_50_ **(C)** conditions from the onset of the steady-state period to pulse force onset (t_0_). The yellow line indicates anticipatory-phase onset (t_A_). The cyan-to-magenta color scale represents normalized firing frequency from low to high values. Dashed circles highlight the peak firing-frequency region of high-threshold motor units (HMU). The horizontal dotted line across (A), (B), (C) stand for 100% firing frequency. The diagonal dotted line extending from the high-threshold to the low-threshold region indicate the 50% firing-frequency, whereas the diagonal dotted line extending across time indicates the recruitment time of medium-threshold motor units.

**Fig. 4.**
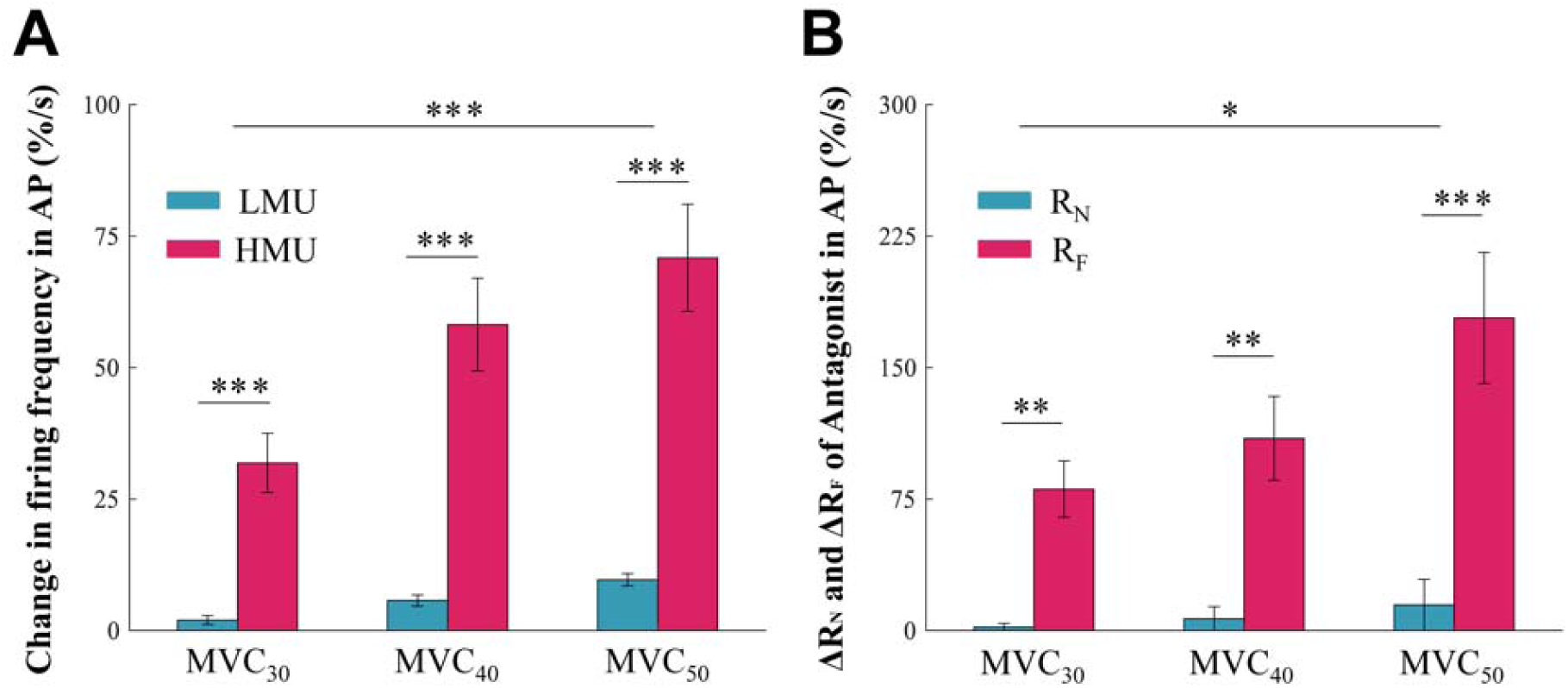
Comparisons of changes in mean firing-frequency of lower-recruitment-threshold motor units (LMU) and higher-recruitment-threshold motor units (HMU) of the agonist muscle during the anticipatory phase (AP) **(A)**. Comparisons between antagonist motor-unit regulation changes in recruited number (ΔR_N_) and firing frequency (ΔR_F_) during AP **(B)**. Asterisks indicate statistically significant difference: * *p* < 0.05, ** *p* < 0.01, *** *p* < 0.001

**Table 2.**
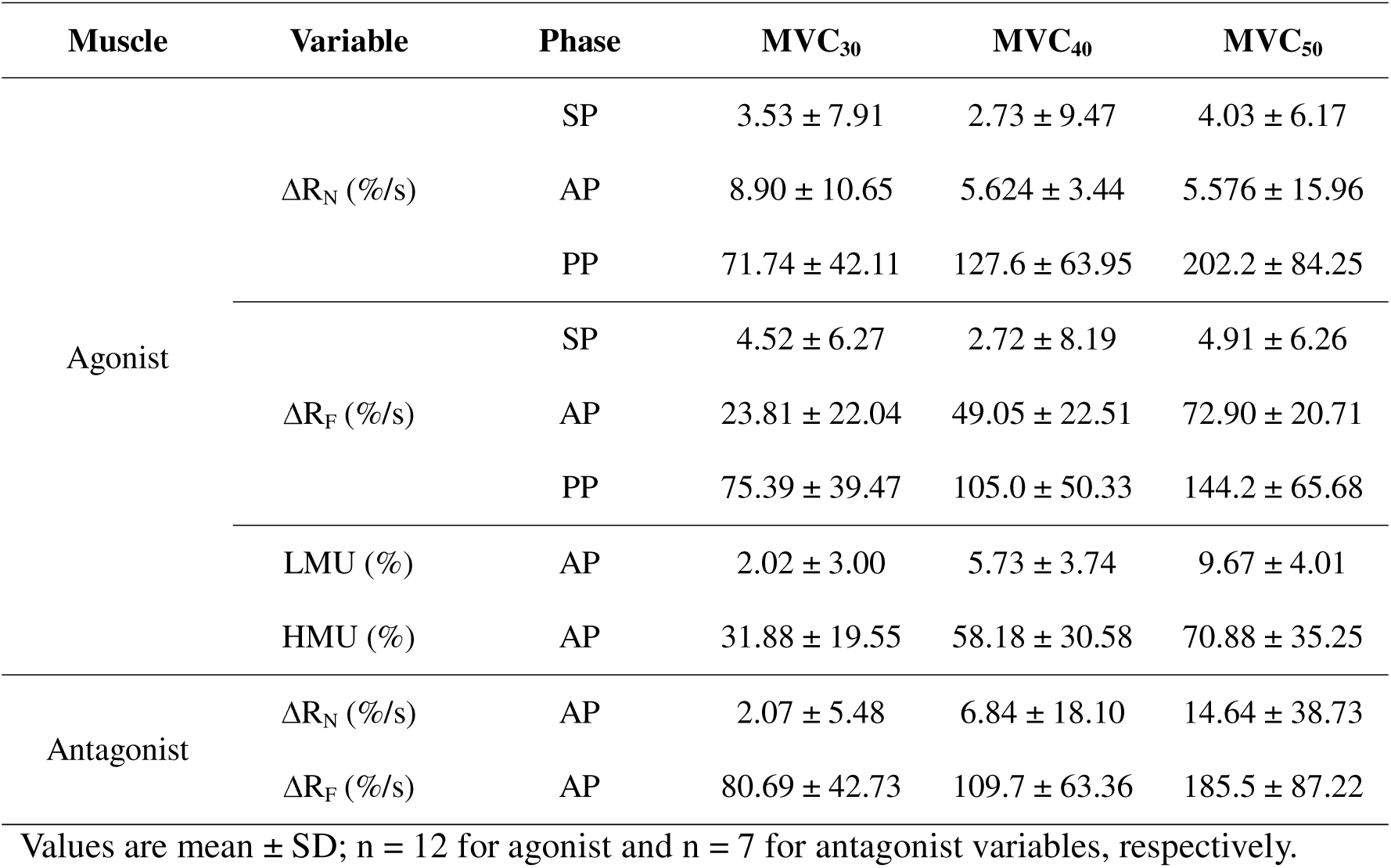
ΔR_N_, ΔR_F_, LMU, and HMU in agonist and antagonist muscles across phases and target force levels.

## 4. Discussion

The primary finding of the study was that anticipatory modulation preceding self-paced pulse force production was predominantly expressed through alterations in motor-unit discharge rate, rather than alterations in the number of detected active motor units. A similar discharge-rate-dominant pattern was observed in antagonist recordings with sufficient decomposition yield, but these results should be considered exploratory. This finding is consistent with preparatory agonist-antagonist coactivation, although it should be interpreted as secondary because antagonist data were available from fewer participants.

The current results showed that the time of feedforward adjustment (t_A_) was approximately 300–350 ms before visible changes in net joint torque by force (Figs. 2A–C). In particular, t_A_ values were similar across force-magnitude conditions (i.e., required net joint torque). This finding is consistent with previous claims that the timing factor in the mechanism of feedforward adjustment is not related to expected perturbation magnitude, that is, perturbation-magnitude independence in time (Olafsdottir et al., 2008; Park et al., 2015; Park and Xu, 2017). Notably, motor-unit recruitment order was constrained by the size principle, which is the gold-standard rule for describing recruitment of multiple motor units in a specific order as a function of force demands. The follow-up question for interpreting the current results is why the controller adjusts the firing frequency of the detected active motor units simultaneously in both agonist and antagonist muscles, rather than recruitment or dropout of motor units, as a feedforward adjustment to prepare for upcoming actions.

To answer this question, it is necessary to identify the advantage of agonist-antagonist coactivation in terms of discharge frequency for upcoming quick force generation. According to the referent-configuration hypothesis, or equilibrium-point hypothesis (Latash, 2008), two parameters, spatial referent coordination (RC or R-command) and apparent stiffness (k or C-command), are assumed to be hypothetical control variables formulated by the equation ΔF = k × (AC − RC), where AC and RC are actual and referent coordinates, respectively (Feldman, 2011). Since quick pulse force could be produced by a fast change in RC, an increase in stiffness (k) as preparation for quick action would have a beneficial effect on the upcoming mechanical outcome (e.g., rate of force change, ΔF). However, this interpretation may not be generalized, because experimental findings have shown that stiffness and RC depend on initial force level (Latash and Gottlieb, 1991).

The increment in discharge rate was greater in HMU than in LMU, both recruited during steady-state force production at 20% MVC. The current results are in line with previous findings showing that high-threshold motor units yield relatively low firing frequency compared with the low-recruitment-threshold units. The firing frequency of higher-recruitment-threshold motor units is always behind that of lower-recruitment-threshold motor units, even during quick and abrupt increases in net muscle force (Desmedt and Godaux, 1977). An interpretation of the feedforward mechanism based on the present motor-unit-level results could be two-fold. First, slow motor units could contribute to the buildup of nervous activity necessary for the forthcoming movement and, specifically, to the control of the excitability of central neurons that generate a phasic discharge to activate hypothesized fast motor units. Second, the greater anticipatory increase in HMU discharge rate may reflect increased preparatory neural drive to the higher-threshold portion of the active motor-unit pool. This interpretation is consistent with orderly recruitment according to the size principle; however, it should not be interpreted as direct evidence of motor-unit contractile type or motoneuron excitability. The absence of measurements of muscle stiffness, spinal excitability, and contractile properties of individual motor units precludes a comprehensive interpretation of the findings. Consequently, the results of this study delineate a motor-unit-level signature of anticipatory preparation, rather than the complete neural mechanism.

This study has several limitations. First, surface decomposition EMG samples detected motor units but did not represent the entire motor-unit pool. This limitation may be addressed by high-density EMG, which covers a larger area. Second, the sample was limited to young men, which may have introduced bias and limited generalizability to women, older adults, and clinical populations. Third, the number of participants in the antagonist analyses was lower because of reduced decomposition yield. Finally, the study did not directly measure muscle stiffness, motoneuron excitability, or motor-unit contractile properties. Therefore, the mechanistic interpretations presented in the Discussion should be considered provisional.

## 5. Conclusions

Preparation for self-paced rapid isometric elbow-flexion force production was accompanied by anticipatory modulation of motor-unit discharge rate. The number of detected active motor units remained relatively stable during AP, whereas discharge rate increased before force onset, scaled with the upcoming force requirement, and was greater in high-threshold motor units. Antagonist recordings indicated a comparable preparatory coactivation pattern; however, this finding should be regarded as secondary because of the lower decomposition yield. Collectively, these findings suggest that discharge-rate modulation is a predominant motor-unit-level characteristic of anticipatory preparation preceding rapid force production.

## CRediT authorship contribution statement

**Ji-Won Park**: Conceptualization, Investigation, Formal analysis, Validation, Visualization, Writing - original draft, Writing - review and editing. **Sungjune Lee**: Data curation, Methodology, Formal analysis, Validation, Visualization, Writing - original draft. **Yoon-Seok Choi**: Formal analysis, Validation, Visualization, Writing - original draft. **Dawon Park**: Formal analysis, Validation, Visualization, Writing - original draft. **Hyeon Hur**: Formal analysis, Validation, Visualization, Writing - original draft. **Hyun-Soo Kim**: Formal analysis, Validation, Visualization, Writing - original draft. **Jaeho Park**: Formal analysis, Validation, Visualization, Writing - original draft. **Jaebum Park**: Conceptualization, Methodology, Resources, Formal analysis, Funding acquisition, Supervision, Writing - original draft, Writing - review and editing.

## Declaration of generative AI and AI-assisted technologies in the manuscript preparation process

During the preparation of this formatted draft, the authors used ChatGPT for manuscript-formatting and language-editing support. After using this tool, the authors reviewed and edited the content as needed and take full responsibility for the content of the submitted article.

## Data availability

All data and code for the current experiments are available from the corresponding author (Jaebum Park) on reasonable request.

## Abbreviations

AP: anticipatory phase
dEMG: decomposition electromyography
EMG: electromyography
HMU: high-threshold motor unit
LMU: low-threshold motor unit
MVC: maximum voluntary contraction
MUAP: motor unit action potential
PP: pulse phase
RF: normalized mean discharge rate
RN: normalized number of detected active motor units
SP: steady-state phase.

