## Supplementary tables for "Anticipatory modulation of motor unit discharge rate before rapid isometric elbow flexion force production"

| **Table 1.** Detected motor unit number and firing frequency of the agonist muscle across phases in 12 participants | | | | | |
| --- | --- | --- | --- | --- | --- |
| **Participant** | **Variable** | **Phase** | **MVC_30_** | **MVC_40_** | **MVC_50_** |
| 1 | Detected MU number | SP | 20 | 17 | 23 |
|  |  | AP | 21 | 18 | 23 |
|  |  | PP | 22 | 21 | 29 |
|  | Firing frequency (Hz) | SP | 14.49 | 10.81 | 11.34 |
|  |  | AP | 14.65 | 11.62 | 12.64 |
|  |  | PP | 14.90 | 13.32 | 16.36 |
| 2 | Detected MU number | SP | 22 | 20 | 18 |
|  |  | AP | 23 | 21 | 19 |
|  |  | PP | 26 | 28 | 28 |
|  | Firing frequency (Hz) | SP | 11.11 | 9.50 | 8.21 |
|  |  | AP | 11.43 | 11.10 | 10.06 |
|  |  | PP | 13.15 | 14.63 | 15.48 |
| 3 | Detected MU number | SP | 17 | 16 | 16 |
|  |  | AP | 18 | 16 | 16 |
|  |  | PP | 20 | 20 | 21 |
|  | Firing frequency (Hz) | SP | 8.87 | 8.66 | 8.37 |
|  |  | AP | 10.10 | 9.71 | 10.30 |
|  |  | PP | 12.26 | 14.17 | 14.57 |
| 4 | Detected MU number | SP | 23 | 23 | 15 |
|  |  | AP | 23 | 22 | 15 |
|  |  | PP | 25 | 28 | 25 |
|  | Firing frequency (Hz) | SP | 10.70 | 9.52 | 5.57 |
|  |  | AP | 10.49 | 9.85 | 5.65 |
|  |  | PP | 12.33 | 14.47 | 11.63 |
| 5 | Detected MU number | SP | 28 | 21 | 23 |
|  |  | AP | 28 | 22 | 24 |
|  |  | PP | 30 | 28 | 34 |
|  | Firing frequency (Hz) | SP | 15.44 | 13.48 | 10.54 |
|  |  | AP | 16.09 | 14.42 | 12.63 |
|  |  | PP | 16.90 | 17.86 | 18.06 |
| 6 | Detected MU number | SP | 17 | 14 | 16 |
|  |  | AP | 19 | 17 | 18 |
|  |  | PP | 22 | 22 | 26 |
|  | Firing frequency (Hz) | SP | 10.13 | 8.27 | 8.09 |
|  |  | AP | 12.32 | 11.40 | 10.49 |
|  |  | PP | 15.42 | 15.87 | 18.02 |
| 7 | Detected MU number | SP | 13 | 17 | 17 |
|  |  | AP | 14 | 19 | 17 |
|  |  | PP | 16 | 25 | 25 |
|  | Firing frequency (Hz) | SP | 10.50 | 9.46 | 9.80 |
|  |  | AP | 11.81 | 10.83 | 11.64 |
|  |  | PP | 13.08 | 14.77 | 16.55 |
| 8 | Detected MU number | SP | 18 | 15 | 18 |
|  |  | AP | 19 | 15 | 19 |
|  |  | PP | 21 | 23 | 29 |
|  | Firing frequency (Hz) | SP | 9.33 | 7.36 | 8.14 |
|  |  | AP | 9.90 | 8.39 | 9.57 |
|  |  | PP | 12.93 | 13.22 | 16.39 |
| 9 | Detected MU number | SP | 17 | 21 | 20 |
|  |  | AP | 18 | 24 | 22 |
|  |  | PP | 19 | 27 | 26 |
|  | Firing frequency (Hz) | SP | 12.94 | 11.94 | 10.71 |
|  |  | AP | 14.77 | 14.46 | 13.15 |
|  |  | PP | 15.95 | 16.75 | 18.35 |
| 10 | Detected MU number | SP | 11 | 17 | 15 |
|  |  | AP | 11 | 17 | 15 |
|  |  | PP | 12 | 19 | 19 |
|  | Firing frequency (Hz) | SP | 9.78 | 10.98 | 9.29 |
|  |  | AP | 9.82 | 11.83 | 8.73 |
|  |  | PP | 10.69 | 14.18 | 13.75 |
| 11 | Detected MU number | SP | 8 | 9 | 11 |
|  |  | AP | 8 | 9 | 11 |
|  |  | PP | 9 | 12 | 13 |
|  | Firing frequency (Hz) | SP | 8.34 | 6.34 | 8.12 |
|  |  | AP | 9.85 | 7.41 | 9.62 |
|  |  | PP | 11.82 | 10.44 | 12.76 |
| 12 | Detected MU number | SP | 14 | 15 | 11 |
|  |  | AP | 15 | 16 | 12 |
|  |  | PP | 15 | 18 | 16 |
|  | Firing frequency (Hz) | SP | 9.60 | 10.26 | 6.87 |
|  |  | AP | 11.09 | 12.12 | 8.64 |
|  |  | PP | 13.04 | 14.93 | 12.99 |

| **Table 2.** ΔR_N_, ΔR_F_, LMU, and HMU in agonist and antagonist muscles across phases and target force levels | | | | | |
| --- | --- | --- | --- | --- | --- |
| **Muscle** | **Variable** | **Phase** | **MVC_30_** | **MVC_40_** | **MVC_50_** |
| Agonist | ∆R_N_ (%/s) | SP | 3.53 ± 7.91 | 2.73 ± 9.47 | 4.03 ± 6.17 |
|  |  | AP | 8.90 ± 10.65 | 5.624 ± 3.44 | 5.576 ± 15.96 |
|  |  | PP | 71.74 ± 42.11 | 127.6 ± 63.95 | 202.2 ± 84.25 |
|  | ∆R_F_ (%/s) | SP | 4.52 ± 6.27 | 2.72 ± 8.19 | 4.91 ± 6.26 |
|  |  | AP | 23.81 ± 22.04 | 49.05 ± 22.51 | 72.90 ± 20.71 |
|  |  | PP | 75.39 ± 39.47 | 105.0 ± 50.33 | 144.2 ± 65.68 |
|  | LMU (%) | AP | 2.02 ± 3.00 | 5.73 ± 3.74 | 9.67 ± 4.01 |
|  | HMU (%) | AP | 31.88 ± 19.55 | 58.18 ± 30.58 | 70.88 ± 35.25 |
| Antagonist | ∆R_N_ (%/s) | AP | 2.07 ± 5.48 | 6.84 ± 18.10 | 14.64 ± 38.73 |
|  | ∆R_F_ (%/s) | AP | 80.69 ± 42.73 | 109.7 ± 63.36 | 185.5 ± 87.22 |
| Values are mean ± SD; n = 12 for agonist and n = 7 for antagonist variables, respectively. | | | | | |
